# HippoMaps: multiscale cartography of human hippocampal organization

**DOI:** 10.1101/2024.02.23.581734

**Authors:** Jordan DeKraker, Donna Gift Cabalo, Jessica Royer, Alexander Ngo, Ali R. Khan, Bradley G. Karat, Oualid Benkarim, Raul Rodriguez-Cruces, Birgit Frauscher, Raluca Pana, Justine Y. Hansen, Bratislav Misic, Sofie L. Valk, Jonathan C. Lau, Matthias Kirschner, Andrea Bernasconi, Neda Bernasconi, Sascha Muenzing, Markus Axer, Katrin Amunts, Alan C. Evans, Boris C. Bernhardt

## Abstract

The hippocampus has a unique microarchitecture, is situated at the nexus of multiple macroscale functional networks, contributes to numerous cognitive as well as affective processes, and is highly susceptible to brain pathology across common disorders. These features make the hippocampus a model to understand how brain structure covaries with function, in both health and disease. Here, we introduce HippoMaps, an open access toolbox and online data warehouse for the mapping and contextualization of subregional hippocampal data in the human brain (http://hippomaps.readthedocs.io). HippoMaps capitalizes on a unified hippocampal unfolding approach as well as shape intrinsic registration capabilities to allow for cross-subject and cross-modal data aggregation. We initialize this repository with an unprecedented combination of hippocampal data spanning 3D *ex-vivo* histology, *ex-vivo* 9.4 Tesla MRI, as well as *in-vivo* structural MRI and resting-state functional MRI (rsfMRI) obtained at 3 and 7 Tesla, together with intracranial encephalography (iEEG) recordings in epilepsy patients. HippoMaps also contains validated tools for spatial map association analysis in the hippocampus that correct for autocorrelation. All code and data are compliant with community standards, and comprehensive online tutorials facilitate broad adoption. Applications of this work span methodologies and modalities, spatial scales, as well as clinical and basic research contexts, and we encourage community feedback and contributions in the spirit of open and iterative scientific resource development.

## Introduction

The hippocampus has long been regarded as a model to understand how brain structure spatially covaries with function (Bahr, 1995; Eichenbaum, 2000). On the one hand, hippocampal anatomy has been recognized to be organized in both anterior-posterior and proximal-distal dimensions (Duvernoy *et al*., 2013; Olsen *et al*., 2019). Anterior-posterior organization is emphasized in foundational descriptions of its anatomical segments (*i.e.,* head, body, and tail) as well as gradual functional differentiation along the hippocampal long axis (Bouffard *et al*., 2023; Poppenk *et al*., 2013; Przeździk *et al*., 2019; Strange *et al*., 2014; Vogel *et al*., 2020; Vos de Wael *et al*., 2018; Palomero-Gallagher *et al*., 2020; Genon *et al*., 2021). Perpendicular to this, there is a preserved arrangement of hippocampal subfields along the proximal-distal (also referred to as medio-lateral) axis (Genon *et al*., 2021; Insausti & Amaral, 2004; Olsen *et al*., 2019; Paquola *et al*., 2020; Ramón y Cajal, 1904; Yushkevich *et al*., 2015; DeKraker *et al*., 2021). These macroanatomical and microstructural features have been suggested to directly relate to hippocampal circuit organization and its embedding within macroscale functional networks (Knierim & Neunuebel, 2016; Leutgeb & Leutgeb, 2007; Rolls, 2016), contributing to specific hippocampal functions and its role as a nexus connecting paralimbic, sensory, and heteromodal association systems, notably the default mode network (Andrews-Hanna *et al*., 2010; Buckner *et al*., 2008; Smallwood *et al*., 2021; Vos de Wael *et al*., 2018). Its broad involvement in multiple networks is clearly compatible with the key role the hippocampus plays in numerous cognitive and affective processes, including memory and language function, together with affective reactivity, stress as well as spatial navigation (Barnett *et al*., 2024; O’Keefe & Nadel, 1978; Stachenfeld *et al*., 2014, 2017; Whittington *et al*., 2022; Cabalo *et al*., 2024). Notably, the hippocampus is also recognized as one of the proximate evolutionary origins of the neocortex (Puelles *et al*., 2019; Sanides, 1969), making it a candidate structure to investigate principles of evolutionary conservation and innovation in the primate lineage (Eichert *et al*., 2023). Collectively, these insights contribute to the notion that the hippocampus is a microcosm of the brain, and that an assessment of its sub-regional organization provides key insights into human neural architectures.

The fine-grained subregional organization of the hippocampus contrasts the somewhat coarse assessment of this structure by most contemporary neuroimaging investigations, which often still treat this complex archicortical structure as a single entity, or even erroneously label it as ‘subcortical’. This is, in part, due to technical limitations: since the hippocampus is thinner and more tightly convoluted than the neocortex, it is difficult to appreciate its cortical architecture in magnetic resonance imaging (MRI) or the extent of its 3D convolutions in sparse histology slices. Relatively few studies have compared its microstructural to mesoscale structural and functional features directly, with most studies opting instead to apply subfield parcellation as a proxy (Caldairou *et al*., 2016; Iglesias *et al*., 2015; Kulaga-Yoskovitz *et al*., 2015; Olsen *et al*., 2019; Romero *et al*., 2017; Yushkevich *et al*., 2010). At the level of the neocortex, there has on the other hand been an increasing repertoire of comprehensive open tools for contextualization of findings, including BALSA (David C. Van Essen *et al*., 2017), NeuroVault (Gorgolewski *et al*., 2015), and NeuroMaps (Markello *et al*., 2022), as well as other contextualization methods incorporated in statistical software such as BrainStat (Lariviere *et al*., 2022) and the ENIGMA toolbox (Lariviere *et al*., 2022) and the multimodal human brain atlas at EBRAINS (https://www.ebrains.eu/tools/human-brain-atlas). With HippoMaps, we now expand anatomy-driven neuroinformatics and multiscale contextualization methods to the human hippocampus.

HippoMaps benefits from multiple recent technical innovations in hippocampal image processing and analysis. First, it leverages a unified hippocampal segmentation and surface mapping approach using deep learning-based image processing (DeKraker *et al*., 2022), imposing a known prior topology (DeKraker *et al*., 2018) and shape-inherent inter-subject alignment (DeKraker *et al*., 2023). Similar to neocortical surface extraction and registration procedures (Boucher *et al*., 2009; Dale *et al*., 1999; Fischl, Sereno, & Dale, 1999; Fischl *et al*., 1999; Kim *et al*., 2005; Lyttelton *et al*., 2007; MacDonald *et al*., 2000), this allows for topology-informed inter-subject registration to a standardized unfolded space (DeKraker *et al*., 2023). This has begun a new wave of high-sensitivity hippocampally-focused studies in topics including the mapping of histology features (DeKraker *et al*., 2020; Paquola *et al*., 2020), blood perfusion (Haast *et al*., 2023; Ngo *et al*., 2023), biophysically-constrained diffusion (Karat *et al*., 2023), hippocampal sclerosis (Ripart *et al*., 2023), neurodevelopmental trajectories (Hanson *et al*., 2023), functional connectivity (Cabalo *et al*., 2023; Lariviere *et al*., 2023; Xie *et al*., 2023), visual receptive field mapping (Leferink *et al*., 2023), and cross-species comparison (Eichert *et al*., 2023). With the increasing aggregation of hippocampal features in a common reference space, it is now possible to devise repositories that allow for a broad contextualization of hippocampal findings. Such work may aid in the interpretation of findings from new studies and experiments, for example by allowing for the cross-referencing of results against established features of hippocampal functional and structural organization.

HippoMaps is conceptualized as an open access toolbox and online data warehouse for hippocampal analysis and multi-scale contextualization. HippoMaps aggregates normative hippocampal data obtained from 3D *ex-vivo* histology, high- and ultrahigh field *in-vivo* magnetic resonance imaging (MRI) at 3 and 7 Tesla, as well as intracranial electroencephalography (EEG) data for the first time into a common, unfolded coordinate system. Moreover, HippoMaps implements a range of non-parametric statistical tests to evaluate the similarity of standardized surface maps, while controlling for spatial autocorrelation within the hippocampal sheet-like topology. This will provide a statistical foundation for accurate enrichment analysis in the hippocampus (Alexander-Bloch *et al*., 2018; Karat *et al*., 2023; Vos de Wael *et al*., 2020). To facilitate broad adoption and continued development, we made scripts (http://github.io/MICA-MNI/hippomaps) and associated data (https://osf.io/92p34/) openly available, and provide expandable online tutorials and guidelines (http://hippomaps.readthedocs.io).

## Methods

### Datasets

To provide broad coverage of many areas of hippocampal research, we initialize HippoMaps with 30 novel minimally processed but spatially aligned data spanning 3D *ex-vivo* histology, high field *in-vivo* structural as well as resting-state functional MRI (rsfMRI), and intracranial electroencephalography (iEEG). These data originate from open source resources including BigBrain (Amunts *et al*., 2013), AHEAD (Alkemade *et al*., 2022), MICs (Royer *et al*., 2022), PNI (Cabalo *et al*., 2024), the MNI open iEEG atlas (Frauscher *et al*., 2018), and are further supplemented with locally collected data including further healthy structural and functional MRI obtained at 3 Tesla and 7 Tesla, as well as iEEG data obtained in epilepsy patients that also underwent pre-implantation multimodal MRI. See the **Supplementary Materials** for details of each dataset and preprocessing.

### Surface mapping

Data processing details are available in the **Supplementary Methods**. Briefly, minimal preprocessing was applied to each dataset using *micapipe v0.2.0* for structural and functional MRI (Cruces *et al*., 2022) and custom code for other data. Though the processing of each data modality differs, they were each mapped to a standardized folded and unfolded surface space using *HippUnfold v1.3.0* (DeKraker *et al*., 2022). Briefly, this entails tissue type segmentation using a deep UNet neural network, fitting of inner, outer, and midthickness surfaces to hippocampal gray matter, mapping to a standardized unfolded rectangular space, and then registration in unfolded space to a standard, histology-derived generated atlas (DeKraker *et al*., 2023). This standardized space is, thus, made equivalent across all subjects. Notably, despite surface meshes having differing tessellations (**Figure 1A**), they can be interpolated in unfolded space to match microscale features (*e.g.,* 3D reconstructed histological stains) to MRI or *vice versa*, spanning a scale of micrometers to millimeters. Using a similar approach, even sparsely sampled data can be spatially mapped across the hippocampus. In this case, we map the centroids of iEEG channels to their nearest corresponding hippocampal vertices. However, in principle, this could also apply to other sparse (or scattered) data such as tissue punches, other invasive recording devices, small resections, or other irregularly spaced sampling methods. We then map iEEG channel data to all vertices within <5mm of the channel centroid, and average data across all channels from all patients with a weighting proportional to geodesic distance from those vertices. This extrapolation method is more robust than a linear or nearest-neighbour extrapolation, which would be strongly driven by only one or a few nearby vertices with data mapped to them, while also still preserving some spatial preference for data from nearby channels. In all cases left and right hemispheres were averaged to increase signal and since no clear differences were seen between them.

**Figure 1.**
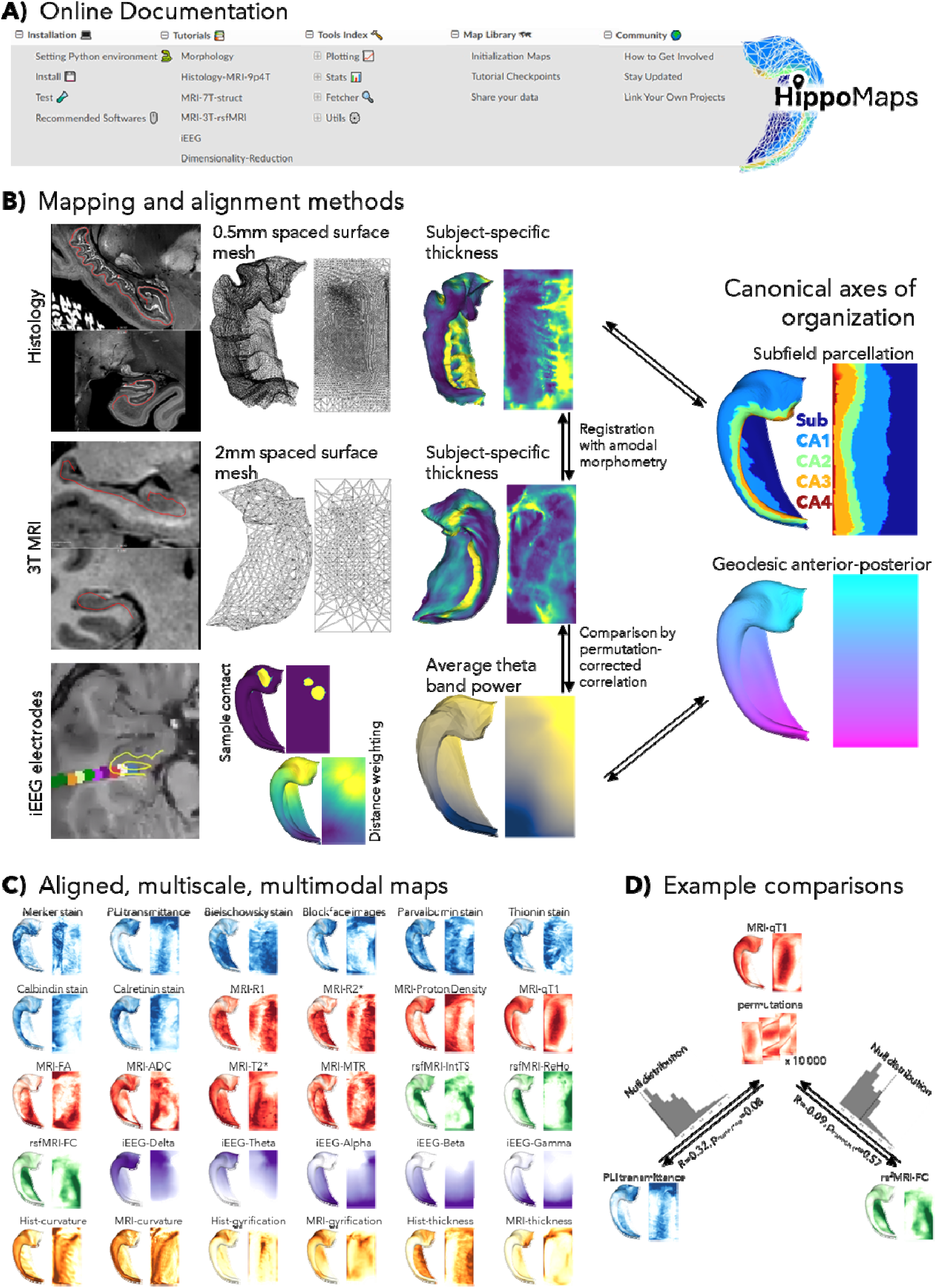
Overview of HippoMaps. **A)** At-a-glance sections of online Documentation. **B)** Surface folding and density are matched to a given sample shape and resolution. Mapping to a standardized unfolded space enables registration and interpolation across scales and data formats, which can then be followed by averaging within a modality, comparison between modalities by spatial correlation, or comparison to anterior-posterior and subfield-related axes of hippocampal organization. **C)** Initial HippoMaps data include 30 highly quality and lightweight surface maps, with extensibility as other experiments are uploaded. Blue maps are derived from histology, red from MRI, green from resting-state functional MRI, violet from iEEG, and yellow from morphology. **D)** Example of how maps are compared via spin test -corrected spatial correlation.

In addition to inner, midthickness, and outer surfaces, any number of intermediate surfaces can be generated at different depths or linearly extrapolated around the outer bounds of the hippocampus (**Figure 2**) (Marcus *et al*., 2011). This is especially useful for sub-millimetric data, where laminar or microstructural profile information can be extracted.

**Figure 2.**
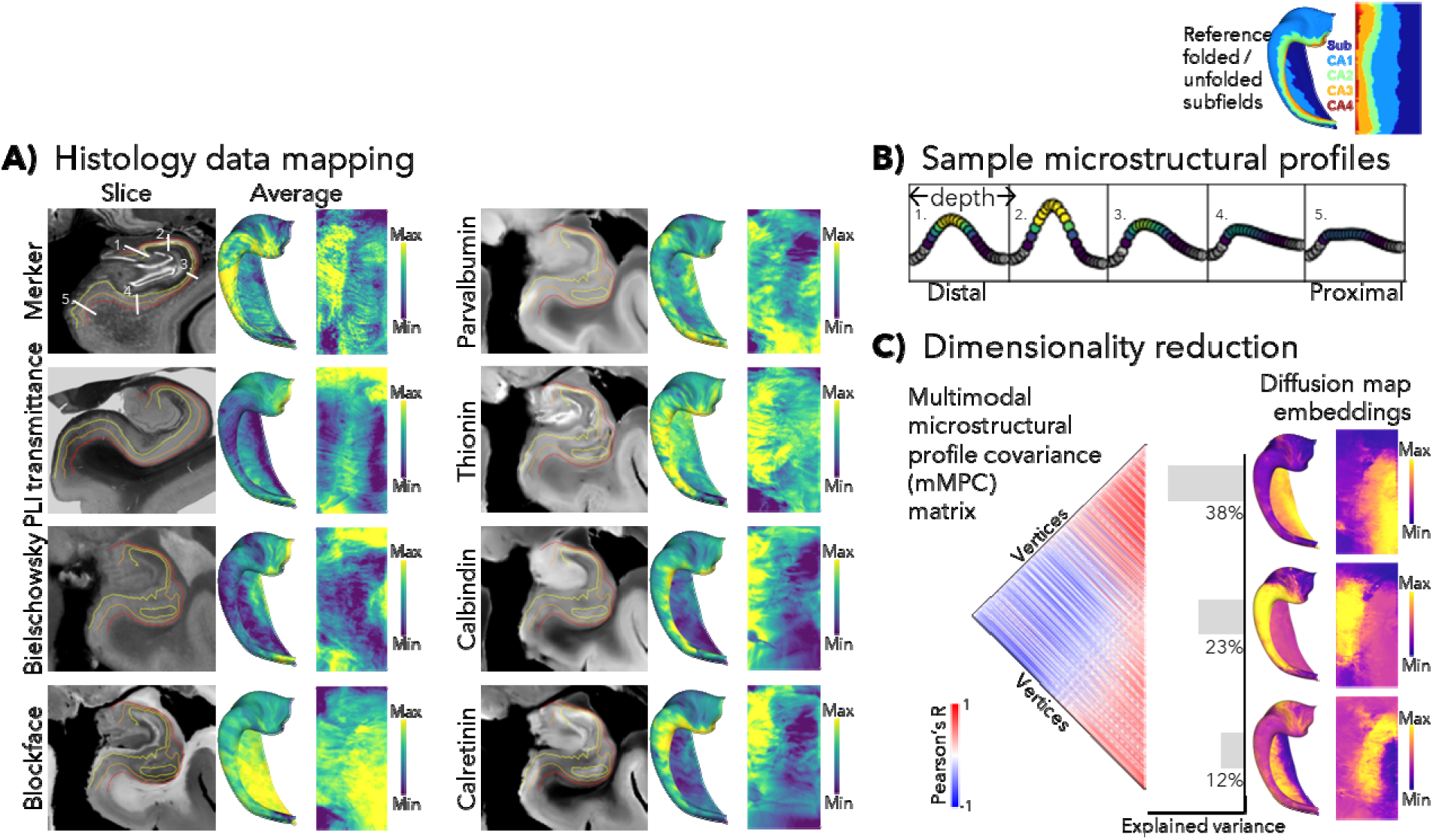
Histology mapping, depth-wise microstructural profiles, and dimensionality reduction. **A)** Sample slices and average 3D maps of histological features. Maps are averaged across depths and, where possible, samples. Numbered lines indicate the approximate locations shown on the corresponding boxes in B). **B)** Example of microstructural profile shapes from five evenly spaced bins across the proximal-distal axis of the BigBrain Merker stain map. Grey indicates points outside of the gray matter mask. **C)** Correlation between microstructural profiles, concatenated across modalities, at each vertex (left). Dimensionality reduction into primary diffusion embedding components 1-3 (right). Scale bars are arbitrary unless indicated otherwise.

### Spatial comparisons

All surfaces have vertex-wise correspondence between subjects and hemispheres, meaning that they can readily be averaged or used in other statistical operations. Similarities between maps can be quantified using spatial correlation, but spatial autocorrelation can compromise significance testing (Alexander-Bloch *et al*., 2018). HippoMaps provides several permutation test to ensure robustness against this issue, including Moran spectral randomization (Wagner & Dray, 2015), “spin” tests (Alexander-Bloch *et al*., 2018; Karat *et al*., 2023; Vos de Wael *et al*., 2020), and “Eigenstrapping” (Koussis *et al*., 2024). **Figure 1D** provides a brief overview of such a correlation using “spin” test permutations. For a detailed overview of the various methods for permutation-corrected spatial correlation, see **Supplementary Methods Figure S2**.

### Dimensionality reduction

Within each method (*i.e.,* histology, structural MRI, resting-state functional MRI, iEEG), we performed dimensionality reduction to summarize the information contained across all group-averaged maps within that methodology. This was also repeated across all maps from all methods (**Figure 7**). Dimensionality reduction consisted of non-linear diffusion map embedding using *BrainSpace* (Vos de Wael *et al*., 2020). Default parameters were used in all cases with a maximum of five components (though only the top three are shown), with the following exception: Pearson’s R was used in calculating affinity matrices in all cases, matching the methods used for spatial comparisons of maps as above, and sparsity was set to 0.1 instead of the default 0.9 in the final reduction across all modalities from all methods to better leverage the richness and reliability of the group-averaged maps. In the case of rsfMRI functional connectivity, components were further contextualized by showing their neocortical counterparts by averaging the connectivity of the top *vs*. bottom 25% hippocampal vertices from each component. The same operation was used to show differences in power spectrum density of the top *vs.* bottom 10% of iEEG components.

## Results

We present novel hippocampal maps in a standardized folded and unfolded space for each of the datasets outlined above. This includes 30 distinct group-averaged maps which have been attentively preprocessed and curated. Within each methodology, some interpretation and summarization via dimensionality reduction is offered, and finally we compare all maps across methodologies in the “Feature combinations” section.

### Histology

Histology is considered a neuroanatomical gold standard, and is the basis for most parcellations and descriptions of brain regions (Amunts *et al*., 2020; Brodmann, 1909; Eickhoff *et al*., 2018; Paquola *et al*., 2019). Here we examined cytoarchitectonic data collected from BigBrain Merker staining for cell bodies (Amunts *et al*., 2013), 3D polarized light imaging (PLI) of neural processes (Axer *et al*., 2011), and the AHEAD dataset with different stains serving as markers of neurons, myelin, and subtypes of interneurons (Alkemade *et al*., 2022) (**Figure 2A**). Most features showed banding in the proximal-distal direction, in alignment with the subfield atlas.

Microstructural (or laminar) profiles are shown for five ROIs across the proximal-distal axis of th BigBrain Merker stain (**Figure 2B**). This is a common method for characterizing laminar structure (Schleicher *et al*., 1999). They show a tight unimodal distribution in the distal CA fields, and a more bimodal distribution in the subiculum as expected based on their known laminar architectures (Duvernoy *et al*., 2013). Profiles for all vertices were concatenated across all stains to make multimodal profiles. That is, for a given vertex, vectors of laminar profiles were concatenated across all modalities, and an affinity matrix was calculated as the correlation between all these extended, or multimodal, profiles, which we call a multimodal microstructural profile covariance matrix (mMPC matrix) (**Figure 2C**). Diffusion map embedding, a non-linear dimensionality reduction technique (Coifman *et al*., 2005; Margulies *et al*., 2016; Vos de Wael *et al*., 2020), decomposed the mMPC matrix into primary components that highlighted the differences between vertices with respect to all modalities and depths. In the first component, a sharp boundary was seen between the subicular complex and proximal CA1 and the rest of the hippocampus. The second and third components in turn highlighted the CA2-3 regions and CA1 with parts of the subiculum, respectively. This is data-driven evidence that subfields across the proximal-distal extent of the hippocampus, rather than anterior-posterior or other patterns, account for structural variance in the hippocampus with respect to these stains. These data-driven decompositions, thereby, echo classical and recent neuroanatomy descriptions of hippocampal microstructure (Ding & Van Hoesen, 2015; Duvernoy *et al*., 2013; Olsen *et al*., 2019).

### Structural MRI

MRI is a key tool for studying human neuroanatomy and structure-function relations due to its non-invasive nature and potential for biomarker discovery. The aggregated *in-vivo* 7 Tesla (7T) and *ex-vivo* 9.4T scanning are especially powerful, achieving greater resolution and contrast than typical 3T or 1.5T clinical scans (Duyn, 2012; Opheim *et al*., 2021). Here, we provide healthy normative maps for such scans (**Figure 3A**) including popular acquisitions: quantitative T1 relaxometry (qT1) and its non-quantitative *ex-vivo* inverse: R1, T2* and its inverse R2*, proton density, diffusion weighted imaging (DWI) estimates of fractional anisotropy (FA) and apparent diffusivity coefficient (ADC), and magnetic transfer ratio (MTR). Note that DWI and MTR images were prone to image artifacts including ringing, leading to ripple-like patterns on individual subject maps and lower inter-subject consistency. Fortunately, since these artifacts are not in-phase between subjects, they are not present in the group-averaged maps.

**Figure 3.**
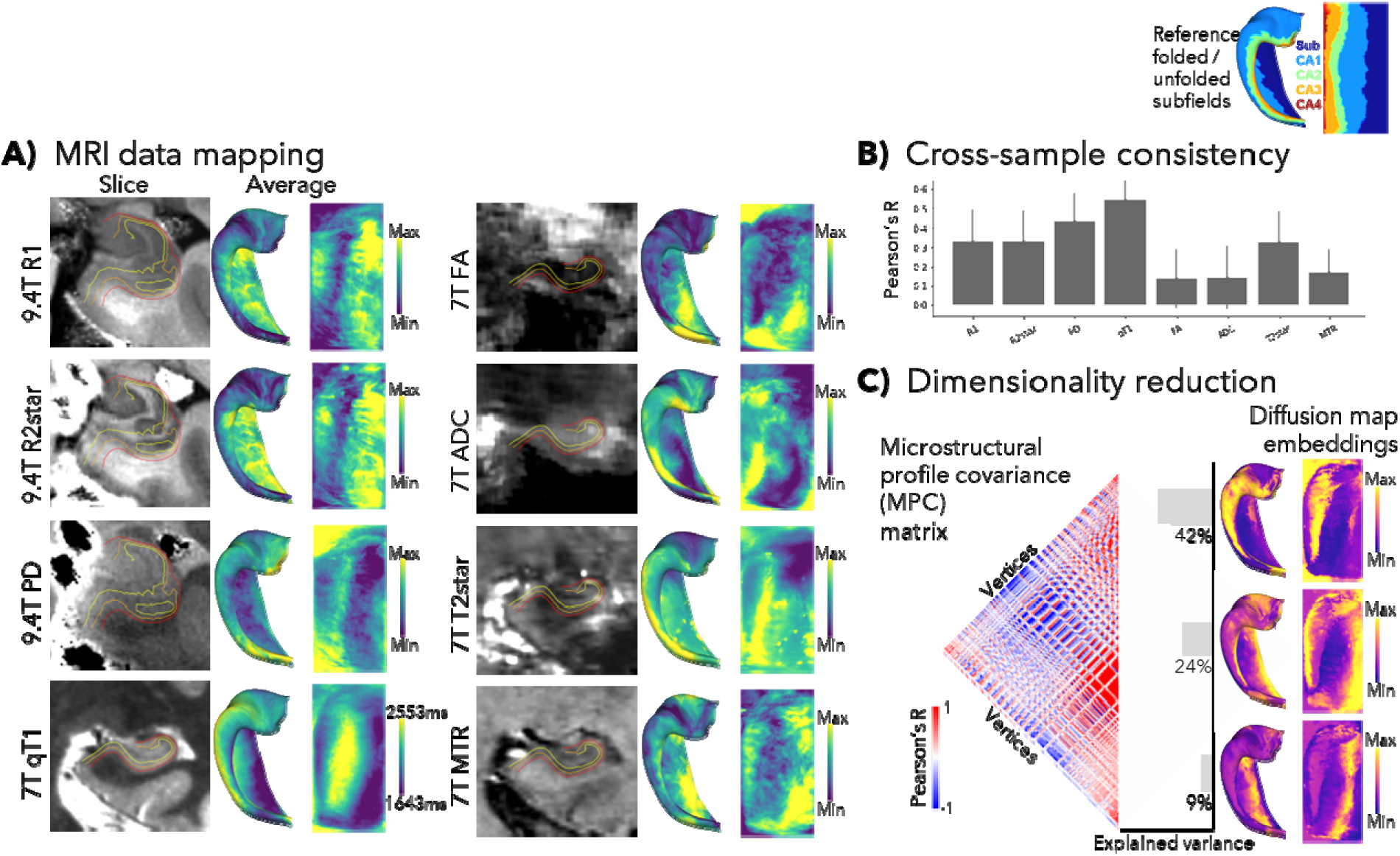
Structural MRI mapping, inter-sample consistency, and dimensionality reduction. **A)** Sample slices and averaged 3D maps of 9.4T *ex-vivo* and 7T *in-vivo* structural MRI features. Maps are averaged across depths and, where possible, samples. **B)** Consistency, as measured by the correlation between all pairs of individual sample maps, **C)** Correlation between microstructural profiles, concatenated across modalities, at each vertex (*left*). Dimensionality reduction into primary diffusion embedding gradients 1-3 (*right*). Scale bars are arbitrary unless indicated otherwise.

Multiple scans were available for averaging (n=4 left+right hippocampi at 9.4T and n=20 left+right hippocampi at 7T), enabling a calculation of consistency across samples via Pearson’s R (**Figure 3B**). DWI and qT1 maps were also calculated in a second validation dataset, consisting of 82 locally scanned healthy participants (including the subset from the MICA-MICs dataset) with a 3T scanner, which showed similar patterns (**Supplementary Results Figure S4**). mMPCs were generated as above and were reduced using diffusion map embedding into primary components, which again highlighted differences across subfields. Only the third component showed anterior-posterior differences, largely within the CA1 subfield.

### Resting-state functional MRI

Functional MRI during the resting state (rsfMRI) allows interrogation of intrinsic brain function via the analysis of spontaneous activity and its statistical dependencies, and has become a key technique in the mapping of functional-anatomical systems (Biswal *et al*., 1997; Buckner *et al*., 2008; Smith *et al*., 2009). Here, we examined several features of rsfMRI in 88 healthy participants scanned at 3T. Intrinsic timescale is a measure of the time it takes for the temporal autocorrelation to drop below a threshold (Golesorkhi *et al*., 2021; Wolff *et al*., 2022) (**Figure 4A**). On a functional level, this is thought to be driven in part by recurrent connections that maintain activity patterns on the order of seconds (Fallon *et al*., 2020). Regional homogeneity considers the similarity between adjacent vertices’ time series, which is thought to indicate the extent of horizontal (*i.e.,* between cortical columns) excitatory connectivity (Zang *et al*., 2004) (**Figure 4B**). Finally, macroscale functional connectivity is by far the most popular rsfMRI feature, with many rich properties that have been explored with respect to white matter connections (Damoiseaux & Greicius, 2009; Greicius *et al*., 2009; Honey *et al*., 2009), network properties (Schmittmann *et al*., 2015; van den Heuvel & Sporns, 2013), organizational gradients (Bernhardt *et al*., 2022; Margulies *et al*., 2016; Paquola *et al*., 2019; Park *et al*., 2021), and many other summary metrics. For simplicity, we examined connectivity between all hippocampal vertices and neocortical parcels from the Schaeffer400 parcellation (Schaefer *et al*., 2018) (**Figure 4C**). The consistency of maps was examined as above, and all measures were significantly greater than zero. Repetition of these analyses in a smaller sample of 7T rsfMRI data showed consistent results **(Figure S4**).

**Figure 4.**
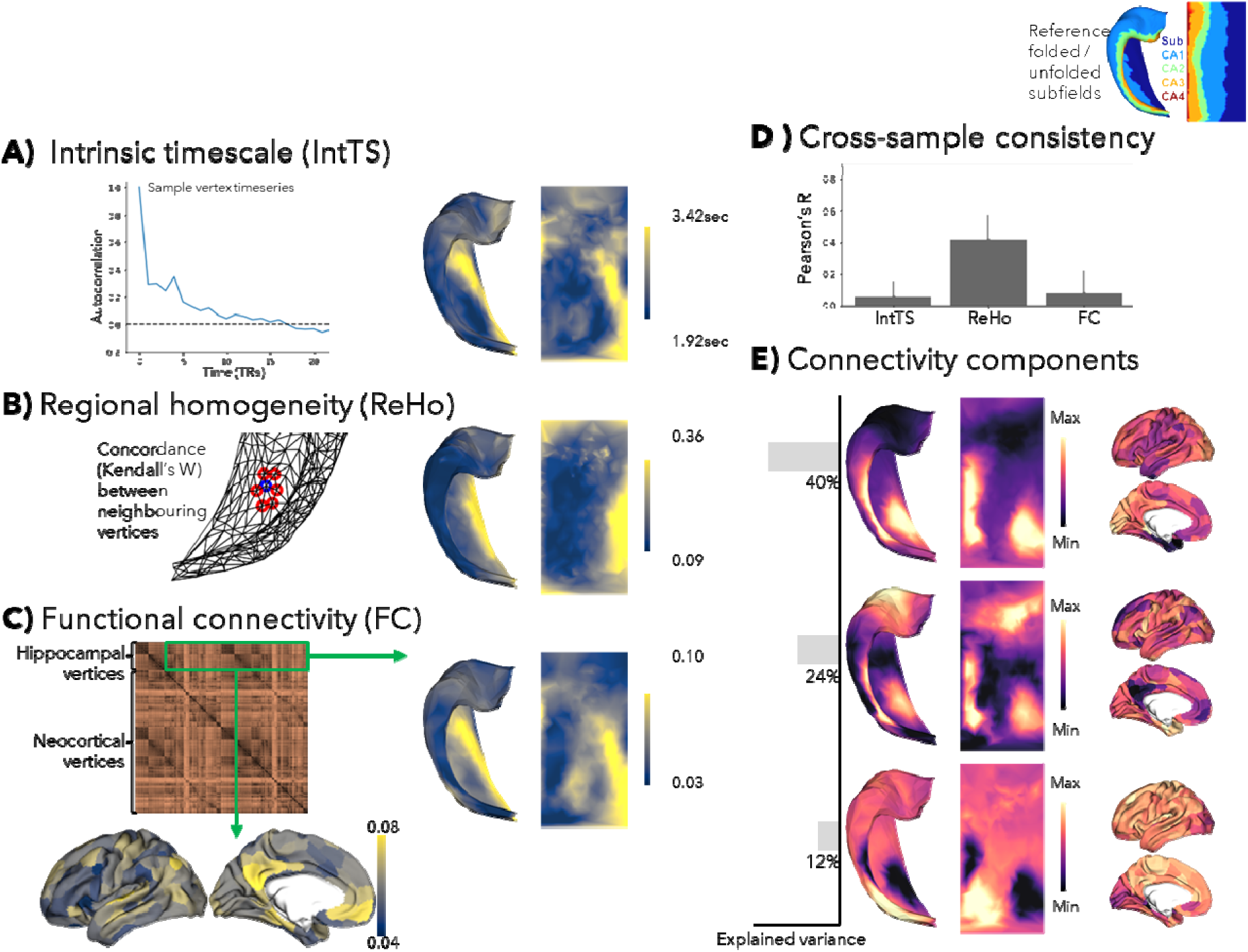
Functional MRI properties. Resting state (rsfMRI) data were used to calculate **A)** intrinsic timescale (recurrence), **B)** regional homogeneity (short range connectivity), and **C)** functional connectivity (long range; to the neocortex). **D)** Cross-sample (that is, subjects and hemispheres) consistency. **E)** Decomposition of functional connectivity patterns across hippocampal vertices into primary diffusion map embedding gradients. Scale bars are arbitrary unless indicated otherwise.

As mentioned above, functional connectivity is a rich measure that can be summarized in many ways. Here, we identified gradual components of intrinsic hippocampal connectivity variations (**Figure 4E**) using the aforementioned non-linear decomposition techniques. Consistent with previous work (Genon *et al*., 2021; Poppenk *et al*., 2013; Przeździk *et al*., 2019; Strange *et al*., 2014; Vogel *et al*., 2020; Vos de Wael *et al*., 2018), we found anterior-posterior differentiation in the first hippocampal component, together with proximal-distal banding with CA1 in particular differing from the other subfields. Neocortical counterparts of this component show that anterior and CA1 regions shared more connectivity with temporal pole, insula, and frontal regions whereas more posterior and non-CA1 subfields shared connectivity with more posterior parietal and visual areas, again consistent with previous findings (Vos de Wael *et al*., 2018). The second component also showed differentiation of CA1 from subiculum and CA2-3 in the more middle and posterior regions, with neocortical correspondences to medial prefrontal and posterior cingulate regions for CA1 and more visual areas for CA2-3 and posterior subiculum.

### Intracranial EEG

Invasive recording methods such as iEEG provide a direct measure of neural activity at high temporal resolution, but typically have lower spatial coverage and are limited to neurological patient populations. In that sense, they can be considered as scattered spatial data, which can be interpolated or extrapolated for hippocampal mapping as described in **Figure 1B**, following previous approaches (Frauscher *et al*., 2018). We employ common measures of the periodic component of iEEG data, as shown by power spectrum density and additionally further simplified to Delta, Theta, Alpha, Beta, and Gamma band powers from low to high frequencies, respectively. Power spectrum densities and band powers derived from hippocampal channels resembled those derived from all channels (**Figure 5A**). Extrapolating channel information across neighbouring vertices from a given hippocampus, a spatial pattern emerged in which both proximal-distal and anterior-posterior differences were seen (**Figure 5B**). Band power is a limited measure of the full power spectrum density though, and so in **Figure 5C** we performed diffusion map embedding of the full power spectrum density. This showed a graded primary anterior-posterior component driven by higher Theta and Alpha power in the posterior and higher Delta power in the anterior hippocampus. The second component showed increased Delta power in the anterior and posterior hippocampus, while the third component showed a slight increase in Delta and decrease in Theta in the subiculum. Results were consistent when using an open iEEG atlas (Frauscher *et al*., 2018) or locally collected data in patients (Paquola, Seidlitz, *et al*., 2020), showing largely conserved patterns in **Figure S5**.

**Figure 5.**
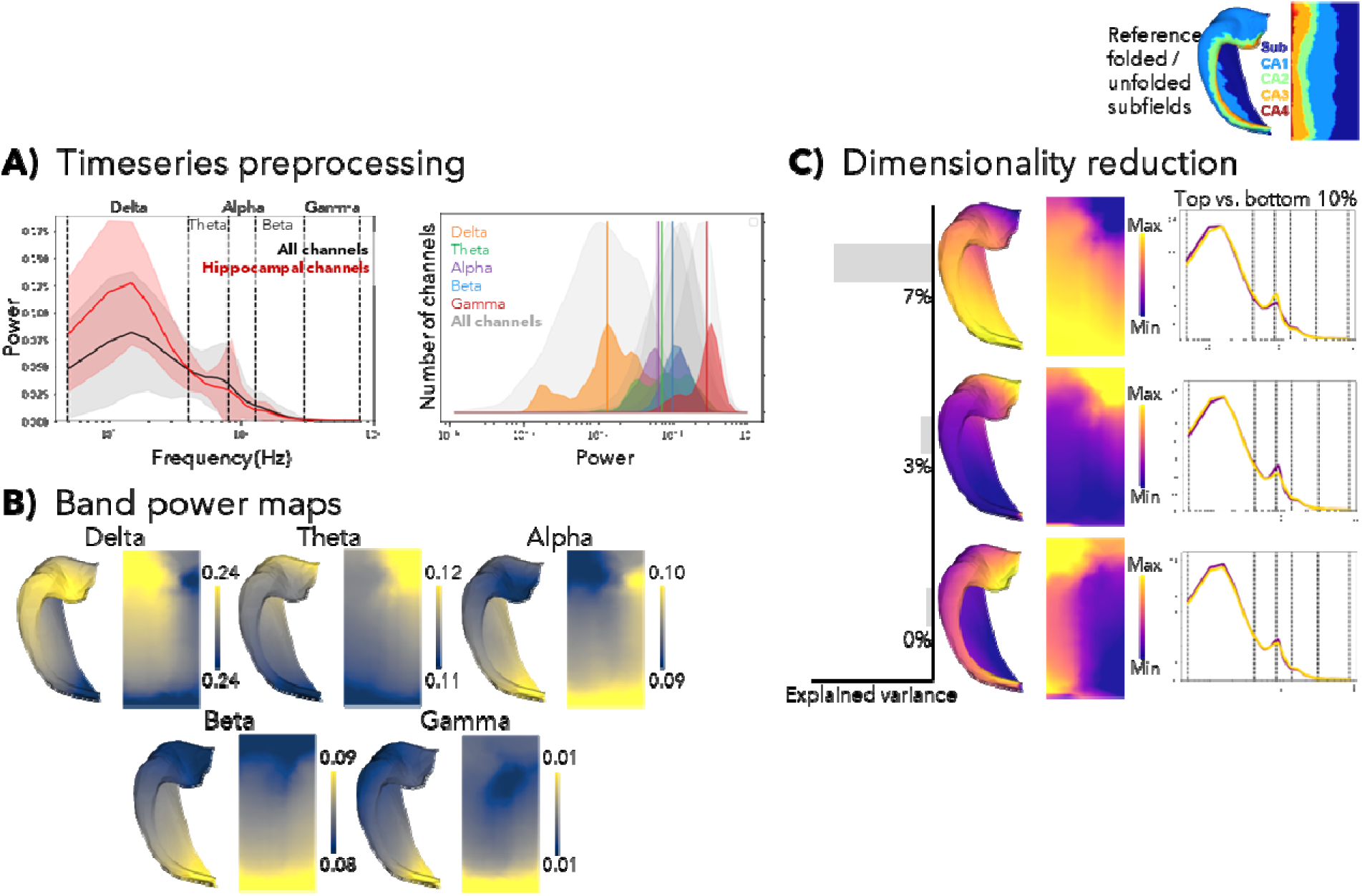
Intracranial EEG (iEEG) properties from time periods deemed “normal” in implanted patients assessed during restin state. **A)** (left) power spectrum density plots of all channels (n=4279) and hippocampal channels (<5mm from any hippocampal midthickness vertex) (n=81), standard deviation shaded. (right) lognormal power within each band for each hippocampal channel, with vertical lines indicating the median and with corresponding bands from all channels in gray. **B)** Spatial extrapolation weighted by geodesic distance shows largely anterior-posterior differences in band powers. **C)** Power spectrum densities reduced into primary diffusion map embedding components. Scale bars are arbitrary unless indicated otherwise.

### Feature combinations

The biggest advantage of a common hippocampal mapping space is that it allows for direct spatial correlation between features from different scales and methods. In **Figure 6A**, we examined relationships between all maps shown above using Pearson’s R with an adapted spin test significance testing to control for spatial autocorrelation in the data (Karat *et al*., 2023). We additionally compared morphological measures of thickness, gyrification, and curvature which are generated within the *HippUnfold* workflow (**Figure S6**). Previous work (DeKraker *et al*., 2020) showed that these features differed between MRI and histology, with the latter showing greater detail including more gyrification and lower thickness. Overall, this revealed many greater-than-chance correlations. This was especially strong within methodologies, but significant correlations between methods employing different spatial scales were also observed.

**Figure 6.**
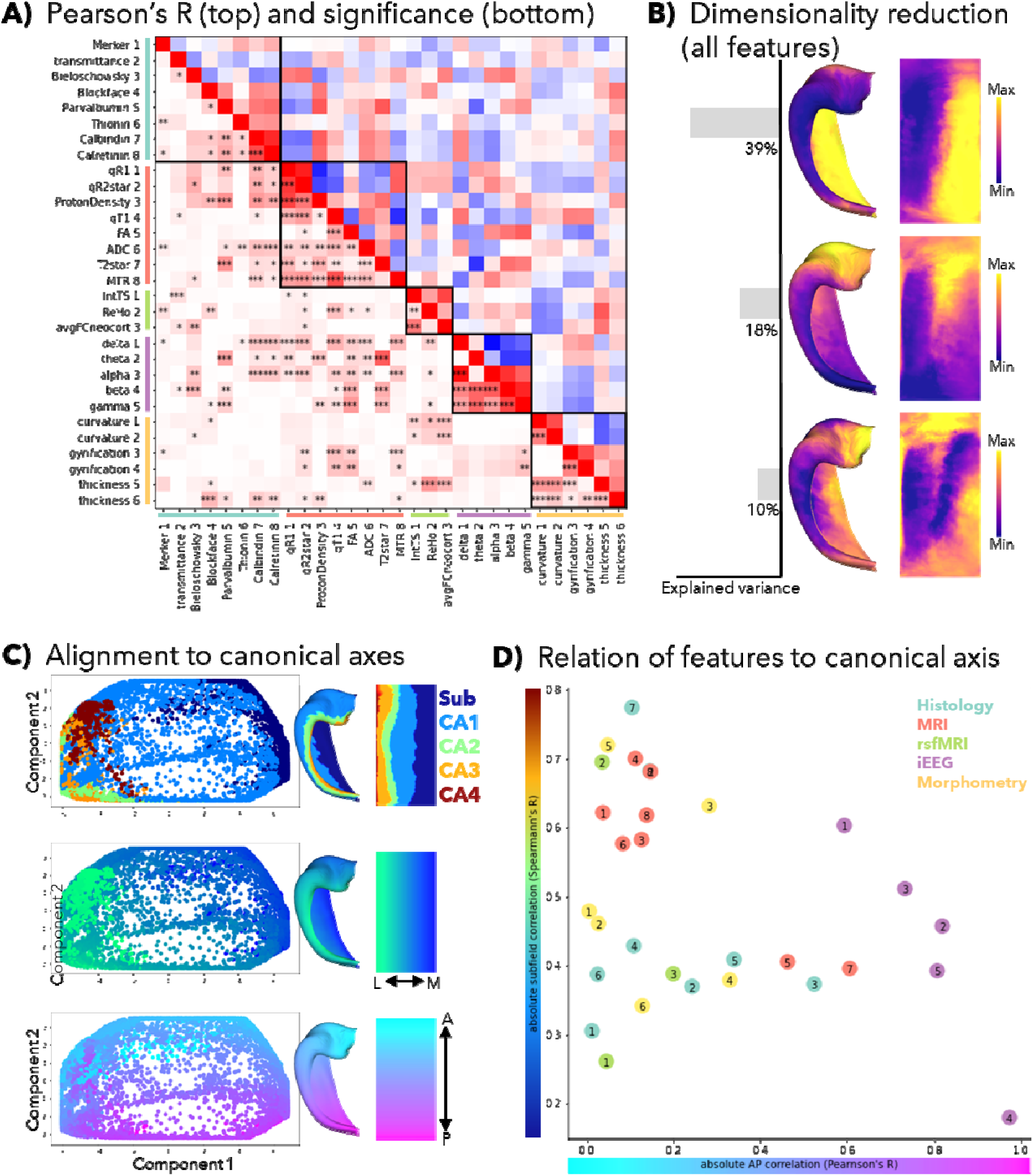
Relationship between all hippocampal maps. **A)** correlation matrix of all features, after resampling to a common 0.5mm vertex-spacing surface. **B)** Diffusion map embeddings 1-3 across all features. **C)** Alignment of gradients 1 and 2 to hippocampal subfields, proximal-distal, and anterior-posterior axes. A = anterior, L = lateral, M = medial, P = posterior. **D)** Absolute correlation between each feature map and the anterior-posterior axis (Pearson’s R) and the maximum permuted subfield labels (Spearman’s R). Scale bars are arbitrary unless indicated otherwise.

We performed a dimensionality reduction as in previous figures, but this time across all features from all methods overviewed here **(Figure 6B)**. As seen in previous results, both proximal-distal or subfield-related and anterior-posterior differences were seen. For additional visualization, we plotted the two most dominant components with colour coding according to subfield and continuous anterior-posterior and proximal-distal gradients (**Figure 6C**). The proximal-distal and anterior-posterior axes of the hippocampus are closely aligned to components 1 and 2, respectively, with component 1 explaining approximately twice the variance (**Figure 6B**). This suggests that while these two axes emerge as natural summaries of many hippocampal feature maps, the proximal-distal direction is stronger.

**Figure 6D** provides a summary of which measures are most correlated with the anterior-posterior and subfield axes of the hippocampus. As expected, the strongest subfield relationships were observed in histological features such as Calbindin and Calretinin staining, or thickness measures at a histological level of precision. Many structural 9.4T and 7T features also showed strong subfield correlations, especially qT1 and qR1. This is encouraging given the increasing availability and adoption of quantitative T1 sequences (Bidhult *et al*., 2016; Haast *et al*., 2016; van der Weijden *et al*., 2021). The employed rsfMRI and iEEG features were still moderately correlated with subfield division, but iEEG and rsfMRI maps showed strong correlations with the anterior-posterior hippocampal axis. Some caution should be exercised here: iEEG data were sparsely sampled and so after extrapolation each band power map was very smooth, which could amplify correlation values (but not significance, since spin test permutations were used to control for spatial autocorrelation). Note also that laminar profiles were not used in this analysis, and histological measures can benefit from the information added by such methods due to their high precision.

### Usability experiment and documentation

HippoMaps as an open toolbox and online data warehouse paves the way for multiple new research avenues, examples of which are shown in **Figure 7**. We anticipate that as hippocampal mapping studies are performed in other research areas, authors can use the initial maps provided here as comparisons and will upload their own maps in the spirit of open and reproducible science, and to boost the visibility of their work. To this end, we provide a set of Python tools, well documented example code to reproduce the maps shown here (labeled as tutorials), and guidelines for how other experimenters should upload their maps to this repository. We have and will continue to answer questions and create community resources via GitHub (https://github.com/MICA-MNI/hippomaps), and all current maps are available on the Open Science Framework (https://osf.io/92p34/).

**Figure 7.**
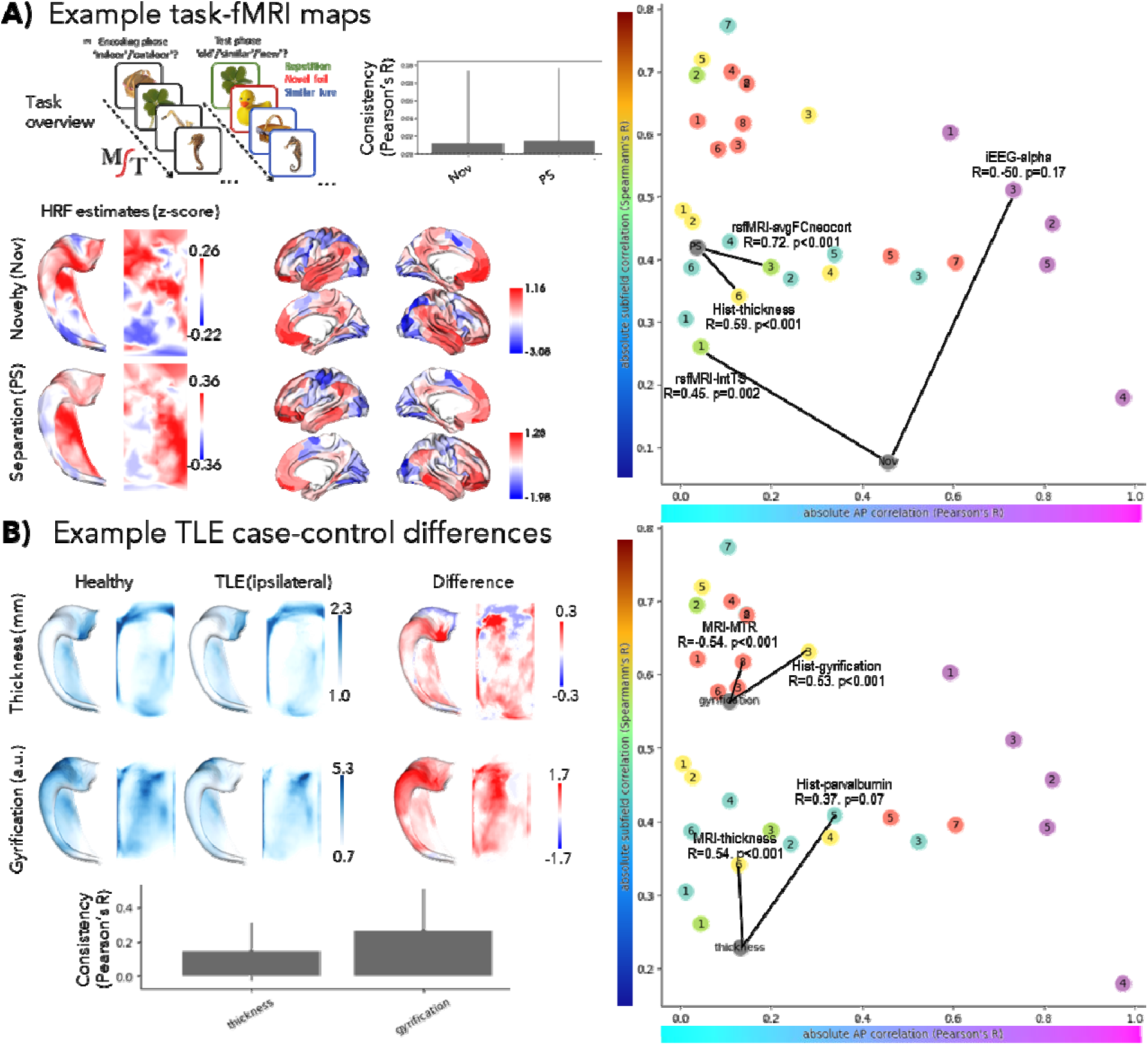
Examples of HippoMaps usage. **A)** Task-fMRI during the Mnemonic Similarity Task (MST) to probe the haemodynamic response function (HRF) magnitudes during successful pattern separation and novel trials. These maps are then compared to all others (right), listing the top two strongest correlations (black lines). **B)** Morphological differences between ipsilateral temporal lobe epilepsy (TLE) patients and healthy controls. Scale bars are arbitrary unless indicated otherwise.

**Figure 7A** illustrates an example experiment with task-fMRI using the Mnemonic Similarity Task (MST) designed to probe pattern separation, a task thought to preferentially involve hippocampal subregions (Pishdadian *et al*., 2020; Stark *et al*., 2019). This can be seen most strongly in subiculum for the successful pattern separation trials, whereas trials with novel stimuli showed anterior-posterior differentiation, similar to previous work (Li *et al*., 2021). Comparing these maps directly to microcircuit features provides context for the demands of these two task conditions: pattern separation was strongly correlated to detailed maps of functional connectivity with the neocortex and histologically-derived thickness, whereas novelty was moderately correlated to iEEG-derived alpha band power and intrinsic timescale (**Figure 7A**, right). Further task-fMRI results from an object-pairing memory task, as well as replication data of the MST at 7T, are shown in **Figure S7**.

**Figure 7B** illustrates an example experiment comparing 33 temporal lobe epilepsy (TLE) patients to 42 healthy, age- and sex-matched controls scanned at 3T MRI. Reductions in hippocampal thickness and gyrification are seen, with the greatest changes in CA1 and CA4 subfields, which have previously been identified as vulnerable areas (Blümcke *et al*., 2012, 2013; Duvernoy *et al*., 2013; Steve *et al*., 2020). Comparing thickness reduction patterns to other maps shows moderate correlations with healthy levels of MRI-derived thickness and parvalbumin staining. Gyrification loss was moderately correlated with MRI-derived magnetic transfer ratios (MTR) and healthy gyrification in histology.

## Discussion

Despite its critical role in human brain organization in both health and disease, the field lacks a standardized framework to aggregate, represent, and compare structural and functional features of the hippocampus. The current work presented HippoMaps as a centralized toolbox and online data warehouse for hippocampal subregional analysis and contextualization. HippoMaps is based on a standardized hippocampal reference space for data aggregation, sharing, and analysis, which leverages recent advances in automated hippocampal segmentation and computational unfolding (DeKraker *et al*., 2022), as well as improvements for cross-modal and cross-subject alignment (DeKraker *et al*., 2023). This repository is initialized with 30 novel maps of hippocampal subregional organization, aggregating a broad array of features from 3D *ex-vivo* histology, *ex-vivo* 9.4 Tesla MRI, alongside with *in-vivo* structural and resting-state functional MRI (rsfMRI) obtained at 3 and 7 Tesla, as well as intracranial encephalography (iEEG) collected from a large cohort of epilepsy patients. This is further extended by a host of tools for visualization and contextualization, as well as online tutorials that recreate the maps shown here and demonstrate how new data can be incorporated and analyzed. HippoMaps will provide key guidance to: *(i)* compare hippocampal features derived from different methods, in particular to cross-reference *in-vivo* imaging measures with *ex-vivo*, *(ii)* interrogate structure-function relationships, for example by contextualizing task-based fMRI findings or intracranial neural recording against spatial patterns obtained from anatomical and microstructural measures, *(iii)* contextualizing case-control deviations in clinical populations against established principles of subregional hippocampal organization, and *(iv)* refining our understanding of hippocampal circuitry, by mapping its functional connectivity and microstructure for a better understanding of its computational operations and transfer functions at the subregional level. HippoMaps is fully open access and designed according to community standards (http://hippomaps.readthedocs.io), to facilitate its dissemination and usability. As such, we anticipate that HippoMaps will represent a powerful analytical ally for fundamental and clinical neuroscientists alike. Considering the unique role the hippocampus plays in human neuroanatomy and cognition (Duvernoy *et al*., 2013; O’Keefe & Nadel, 1978) and its important computational properties (Knierim & Neunuebel, 2016; Leutgeb & Leutgeb, 2007), it may furthermore provide key insights and guidance into the design and validation of emerging bio-inspired AI architectures.

We anticipate that surface-based registration will become the standard for subregional hippocampal mapping, as it has in the neocortex (Fischl, *et al*., 1999; Glasser *et al*., 2013; Ma *et al*., 2023; Robinson *et al*., 2014; Van Essen *et al*., 1998). HippoMaps is a major step in advancing the usability of this methodology, generating utilities, scientific context, and an open community for examining the hippocampus in detail. Moreover, our repository is designed to employ the same data standards that have already been extensively developed for neocortical brain imaging data including Brain Imaging Data Standards (BIDS) (Olsen *et al*., 2019; Yushkevich *et al*., 2015); NIfTI/GIfTI file formatting (Glasser *et al*., 2013); and Findability, Accessibility, Interoperability, and Reusability (FAIR) principles (Olsen *et al*., 2019; Yushkevich *et al*., 2015). Our online tutorials also showcase the straightforward interplay between *HippUnfold*, HippoMaps, and other community tools for surface analysis including *Connectome Workbench*, *BrainStat*, and *NiLearn*. Despite its demonstrated benefits, surface-based alignment is not yet universal for the neocortex and certainly still in its infancy for the hippocampus. Thus, while we encourage the use of surface-based methods, we also provide code and examples of how to map volumetrically aligned hippocampal data (*e.g.,* in a standard volumetric space such as MNI152 or others) to hippocampal surfaces for comparison and contribution to HippoMaps. In the field, work progresses at the level of hippocampal subfield parcellation at the level of histology, for example to derive additional subregional divisions (González-Arnay *et al*., 2024; Henriksen *et al*., 2010; Igarashi *et al*., 2014). Moreover, there have been ongoing efforts by the neuroimaging community to harmonize boundary heuristics (Olsen *et al*., 2019; Yushkevich *et al*., 2015). Under the HippoMaps framework, descriptions go beyond typical unitary descriptions of the hippocampus and beyond its parcellation into subfields to the level of mapping vertex-wise or columnar structure of hippocampal archicortex. The columnar level represents an important structural and functional modularization of the brain (Mountcastle, 1997), and has the potential to unlock new facets of hippocampal computation. As such, different subfield parcellations can also be converted to surface format and integrated seamlessly within the HippoMaps warehouse. Thus, we apply considerable futureproofing, and we encourage the broader hippocampal research community to upload their own maps to this repository under our support, curation, and online guidelines and tutorials.

HippoMaps critically depends on the quality of repository data. Some maps varied between individuals, as reflected in lower consistency metrics. This was strongest in diffusion-derived MRI, magnetization transfer, and functional MRI maps. This can reflect idiosyncratic imaging artifacts that average out over large samples, or systematic imaging artifacts like dropout near major tissue interfaces. In fMRI this could also reflect true inter-individual variability that is not accounted for by structural alignment alone. It is notable that some features showed extreme intensity values at the anterior and posterior edges - these are relatively small in native space and so have limited constituent data and are prone to interpolation artifacts. Thus, the anterior and posterior edges of each map should be interpreted with some caution. Consistency was not evaluated in histology due to small sample sizes, and histology is generally less scalable than MRI making averages across many samples costly. However, as new high throughput methods become more widespread, invaluable datasets such as BigBrain, AHEAD, and others, can provide a high level of anatomical precision and a breadth of extracted features. Future data uploaded to HippoMaps should, thus, aim to include state-of-the-art acquisition methods, averages over many samples where possible, and apply robust preprocessing and quality control to minimize artifacts that limit the quality of comparisons and conclusions about hippocampal organization that can be drawn.

Multi-feature aggregation as in the HippoMaps repository provides extensive opportunities to assess relationships between hippocampal structure and function, to cross-validate *in-vivo* measures with *ex-vivo* imaging as well as histological data. Structural and microstructural data derived from 3D histology and MRI currently aggregated support a close alignment of many feature maps with the classic subfields account of the hippocampal circuitry. Moreover, several measures, particularly those derived from functional modalities such as rsfMRI or iEEG, lend additional evidence for anterior-posterior differentiation of the hippocampal formation. Specifically, diffusion map embedding of hippocampal rsfMRI connectivity and iEEG power spectrum densities showed that anterior-posterior differentiation captured most inter-regional variance, whereas histological and structural MRI measures showed primarily proximal-distal or subfield-related differentiation. The consistently repeated structural motifs across the anterior-posterior axis of the hippocampus are suggestive of parallel repeated computations being performed on different input and output information across the anterior-posterior hippocampal axis, in line with prior accounts (Poppenk *et al*., 2013; Strange *et al*., 2014). These two dimensions have also been suggested to topographically represent the functional embedding of the broader mesiotemporal region in large-scale functional networks, in particular default mode and multiple demand networks (Andrews-Hanna, Reidler, Sepulcre, *et al*., 2010; Buckner *et al*., 2008; Duncan, 2010), which provides a potential substrate for the parametric mixing of both functional systems in macroscale brain function (Paquola, *et al*., 2020). It is, therefore, not surprising that two axes explain the greatest proportion of the variance across all maps in the current repository as well, consolidating the notion that a two dimensional organization may serve as a powerful summary descriptor for a broad array of hippocampal structural and functional features (Genon *et al*., 2021).

We provide adapted methods to control for autocorrelation when comparing spatial maps to one another in the hippocampus. We specifically adapted Moran’s spectral randomization,“spin test” permutation, and Eigenstrapping permutation testing that have previously been introduced to study neocortical data (Alexander-Bloch *et al*., 2018; Karat *et al*., 2023; Vos de Wael *et al*., 2020; Wagner & Dray, 2015, Koussis *et al*., 2024). These methods reveal robust correlations between many of the maps included here. Many of these relationships support the validity of the methods being applied, for example between *in-vivo* qT1 and *ex-vivo* R1 which are inverses of one another. Another example is that functional connectivity of the hippocampus was strong to default mode neocortical areas, as shown in previous work (Andrews-Hanna *et al*., 2010; Norman *et al*., 2021; Vos de Wael *et al*., 2018; Ward *et al*., 2014), with connectivity being strongest in the subiculum. This recapitulates the role of the subiculum as the primary output structure of the hippocampus, and contributions of the hippocampus to functions typically ascribed to the default mode network such as mind-wandering, episodic recall, or future-thinking that are frequent during rest (Bellana *et al*., 2017; Buckner, 2010; Christoff *et al*., 2016; Fox *et al*., 2015; Ross & Easton, 2022; Schacter *et al*., 2017; Yang *et al*., 2020). Some relationships reveal novel information about the methods themselves: PLI transmittance is thought to reflect many microscopic structures under the broad heading of “neural processes” or “nerve fibers” (Axer *et al*., 2001; Dammers *et al*., 2012). Across the extent of the hippocampus, this feature correlated with Bielschowsky and Thionin staining, R2*, average neocortical functional connectivity, and, most significantly, rsfMRI intrinsic timescale. Intrinsic timescale is hypothesized to relate to recurrent connections (Chaudhuri *et al*., 2014), which could indeed be supported by dense neural processes. Finally, we illustrate contextualization via nonlinear diffusion map embedding across maps. When applied to all maps, we show data-driven separation of subfields, in line with previous work. We also note that in this latent space, CA4 closely resembles CA1, even though they are not adjacent topologically. This fits descriptions of CA4 as having a wide pyramidal layer with large and dispersed neurons, similar to CA1 (Duvernoy *et al*., 2013), and indeed in some cases these two areas have similar disease vulnerabilities, for example in drug-resistant temporal lobe epilepsy (Blümcke *et al*., 2012). Future work may determine more selectively what features make these two regions similarly vulnerable, or explore conditions with differential vulnerability.

At the level of the neocortex, several packages already exist to facilitate the contextualization of results (Larivière *et al*., 2023, 2021; Markello *et al*., 2022). With HippoMaps, such an approach is now also possible for the hippocampal region, and we demonstrate the contextualization of task fMRI maps during an episodic memory paradigm as well morphological alterations in patients with temporal lobe epilepsy relative to healthy individuals. Such approaches can help to clarify the hypothetical role of microstructural features in specific hippocampal computations, such as pattern separation (Bakker *et al*., 2008; Leutgeb *et al*., 2007; Schmidt *et al*., 2012), pattern completion (Guzman *et al*., 2016; Leutgeb & Leutgeb, 2007), and novelty detection (Chen *et al*., 2011; Larkin *et al*., 2014). These previously assumed relations of function to microstructure have generally relied on parcellations of the hippocampus into stereotyped subfields; with HippoMaps, it is instead possible to compare functional and microstructural maps directly without any predefined subfield labeling. In addition to offering potential increases in anatomical specificity, this representation may also lend itself more naturally to sensitive spatial correlation with autocorrelation control through permutation testing. One area for future work will lie in consolidating mesoscale connectivity with detailed descriptions of the internal hippocampal circuitry, which will not only help to further understand the computations of specific hippocampal subregions but which may also clarify the different substrates of computation (Beaujoin *et al*., 2018; Bennett & Stark, 2016; Berron *et al*., 2016; Karat *et al*., 2023; Lacy *et al*., 2011; Ly *et al*., 2020). Indeed, hippocampal circuitry has inspired the basic ways in which we think about biological computation, spurring principles such as long-term potentiation (Hebb, 2005), and carrying important computational models like the Boltzmann machine (Ackley *et al*., 1985) and Tolman Eichenbaum machine (Whittington *et al*., 2020). Even more recent computational models and associated theory still center around hippocampal structure as told through a stereotyped subfield architecture (Gandolfi *et al*., 2023; Whittington *et al*., 2020). Formal mapping, rather than stereotyped descriptions, can extend this work, building up biological plausibility of such models and scaffolding our understanding of these systems. For this reason, HippoMaps may also provide precise macro-, meso- and micro-scale hippocampal features in a common same space to further identify and harness computational properties of its circuitry.

## Ethics & Inclusion

The data used in HippoMaps was sourced from multiple open datasets. The MICs dataset was obtained with the approval of the Ethics Committee of the Montreal Neurological Institute and Hospital (2018– 3469). The PNI dataset was obtained with approval by the Research Ethics Board of McGill University. The iEEG dataset was obtained with approval from the MNI as lead ethics organization (REB vote: MUHC-15-950). All participants from the MICs, PNI, and iEEG datasets provided written informed consent, which included a provision for openly sharing all data in anonymized form. The AHEAD dataset samples were collected from the body donation program of the University of Maastricht following a whole-body perfusion, for which written consent was obtained during life, and in accordance with the Dutch Burial and Cremation Act. The 3D-PLI dataset was collected through the body donor program of the University of Rostock, Germany, and in accordance with the local ethics committee. The BigBrain dataset was collected through the body donor program of the University of Düsseldorf in accordance with legal requirements. No further ethics approval was needed for the present study.

All listed authors contributed to either conception and design of the project, data acquisition, curation, analysis, interpretation, software development, or manuscript preparation in accordance with Nature Methods’ authorship criteria. Author roles and responsibilities were additionally defined during regular meetings of the Helmholtz International BigBrain Analytics and Learning Laboratory (HIBALL), which focuses on integrating and sharing multiscale and multimodal neuroimaging data.

## Supporting information

Supplementary Results

## Acknowledgements

JD was supported by a Natural Science and Engineering Research Council of Canada Post Doctoral Fellowship award (NSERC-PDF), and the Helmholtz International BigBrain Analytics and Learning Laboratory (HIBALL), supported by the Helmholtz Association’s Initiative and Networking Fund and the Healthy Brains, Healthy Lives initiative at McGill University

BK was supported by a NSERC postgraduate scholarship.

JR was supported by a Canadian Institute of Health Research Canada (CIHR) fellowship award.

DGC was supported by Fonds de recherche du Quebec-Sante (FRQS) and Savoy Foundation Doctoral fellowship award.

RCR acknowledges that this research was undertaken thanks in part to funding from the Ministère de l’Economie, de l’Innovation et de l’Énergie du Québec and to funding from the Healthy Brains, Healthy Lives (HBHL) initiative at McGill University, and FRQS.

KA received funding from the European Union’s Horizon Europe Programme, grant agreement 101147319 (EBRAINS 2.0 Project, KA, NPG).

MA was supported by computing time and was granted through VSR Computing Time Projects on the supercomputer JURECA at JuLlich Supercomputing Centre (JSC), Germany, to analyze and 3D reconstruct the 3D-PLI data set.

BF was supported by a project grant from CIHR (PJT-175056)

MK received funding from the Swiss National Science Foundation (SNF_219240) SLV is supported by HIBALL and the Jacobs foundation.

JCL was supported by an NSERC Discovery Grant (RGPIN-2023-05562), research start-up funding from the Department of Clinical Neurological Sciences at Western University, and a McGill-Western Initiative for Translational Neuroscience (ITN) Impact Grant.

BCB acknowledges research support from NSERC (Discovery-1304413), CIHR (FDN-154298, PJT-174995, PJT-191853), SickKids Foundation (NI17-039), HIBALL, HBHL, Brain Canada Foundation, the Montreal Neurological Institute, and the Tier-2 Canada Research Chairs Program.

## Notes

https://hippomaps.readthedocs.io/en/latest/

https://github.com/jordandekraker/hippomaps

https://osf.io/92p34/

### Competing Interest Statement

The authors have declared no competing interest.

### Summary of Updates

Additional methods details, fully operable online Tutorials, and rework of Introduction and Figure 1 emphasizing applications, as per Reviewer suggestions

