## Supplementary Results for "HippoMaps: multiscale cartography of human hippocampal organization"

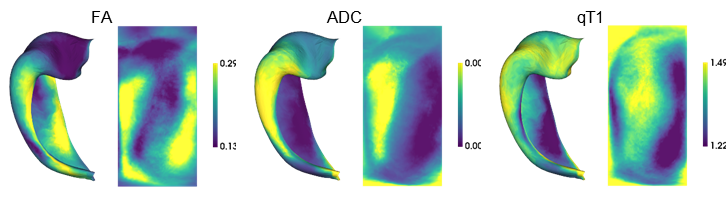

**Figure S3.** Replication of a subset of structural MRI features from **Figure 3** in an independent, 3T scanned cohort (n=82).

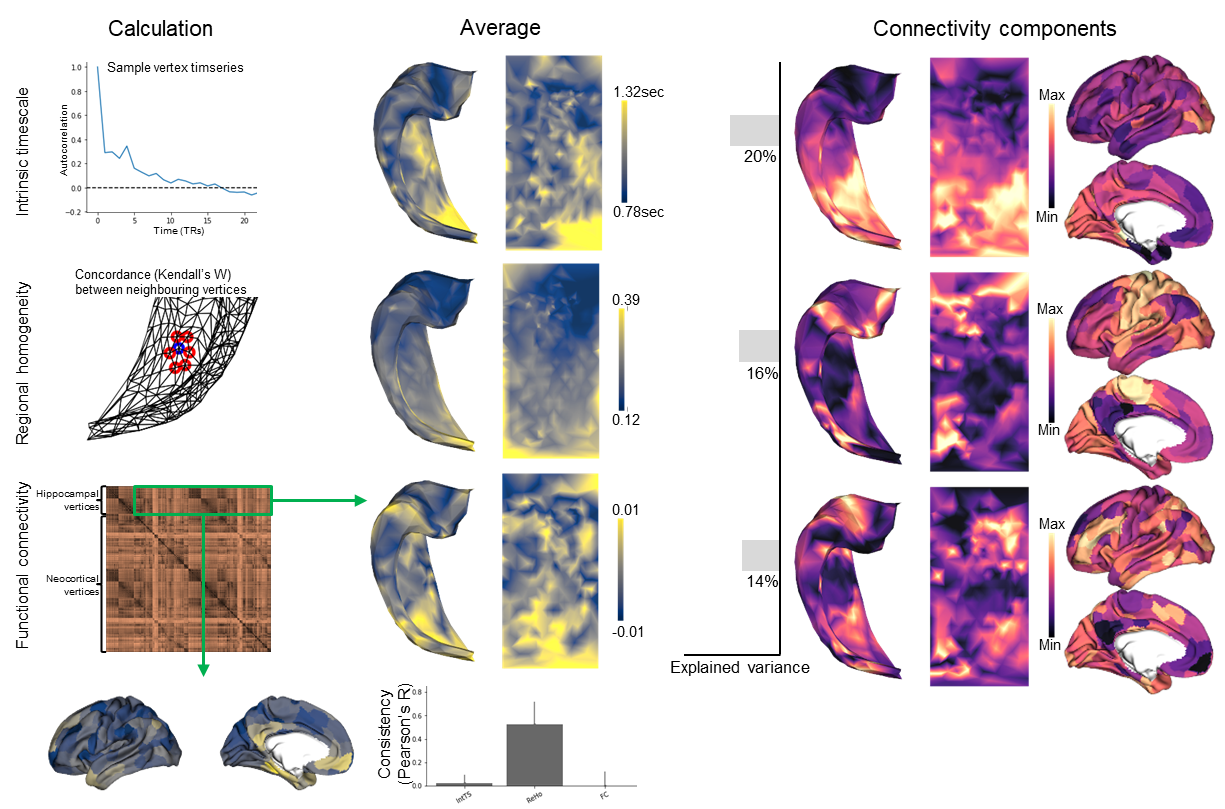

**Figure S4.** Replication of functional MRI properties from **Figure 4** in an independent 7T cohort (n=10).

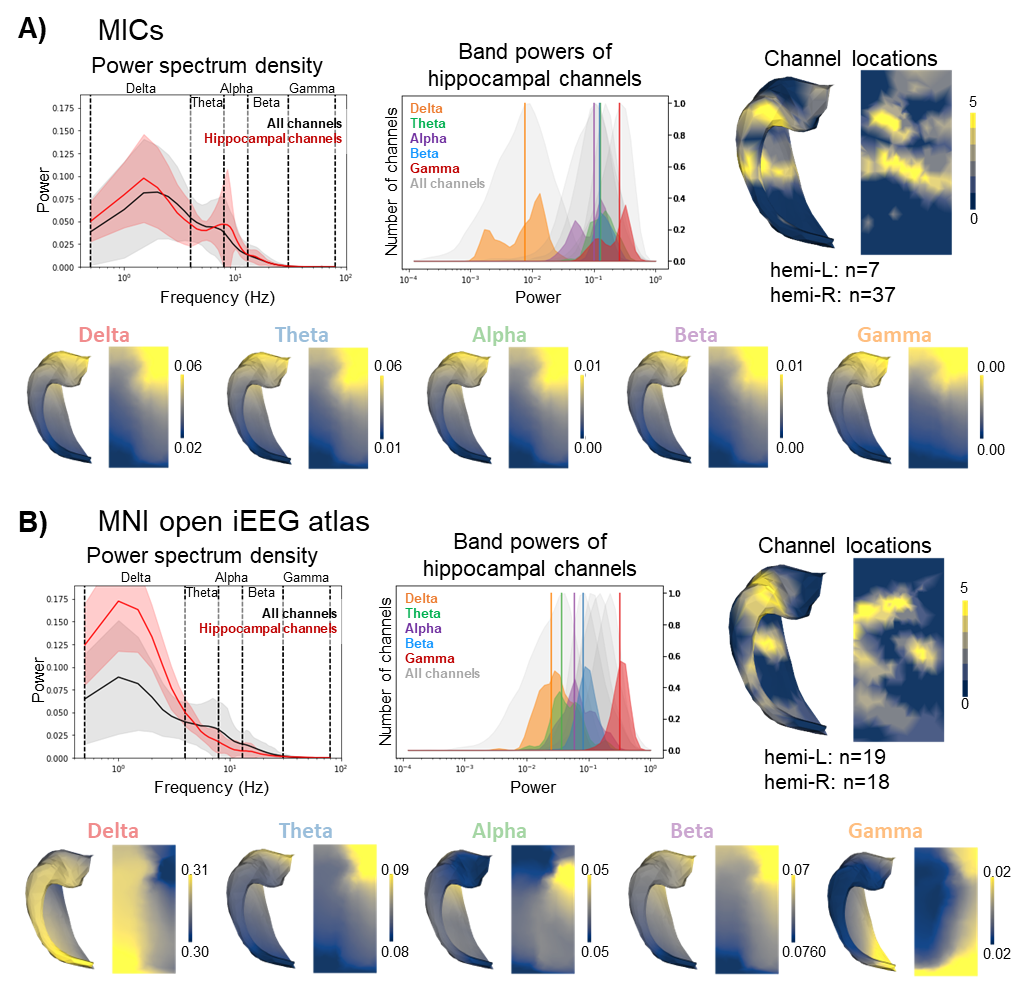

**Figure S5.** Separation of iEEG properties in **Figure 5** into component datasets for replication. **A)** Locally collected iEEG, resampled to 200Hz to match the Frauscher *et al.* (2018) open source dataset. **B)** Open source dataset analysis based on Frauscher *et al.*

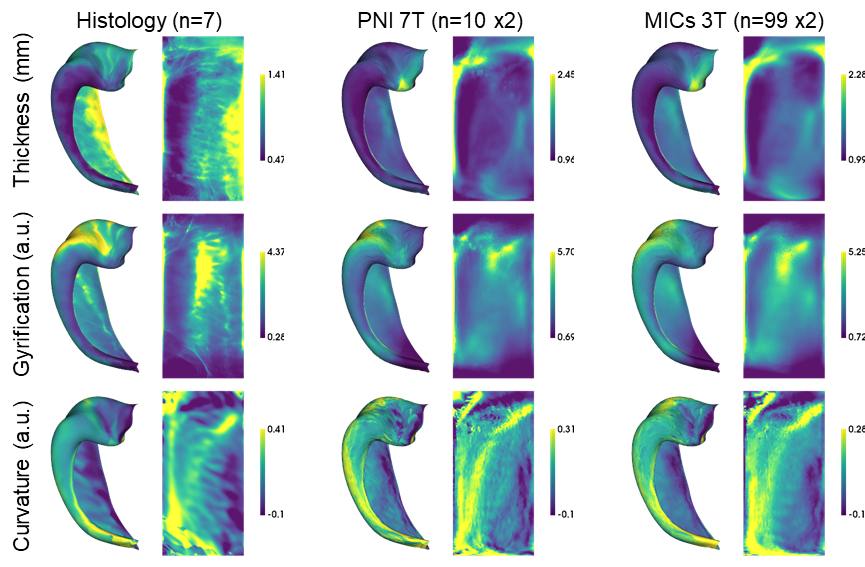

**Figure S6.** Morphometric thickness, gyrification, and curvature across datasets.

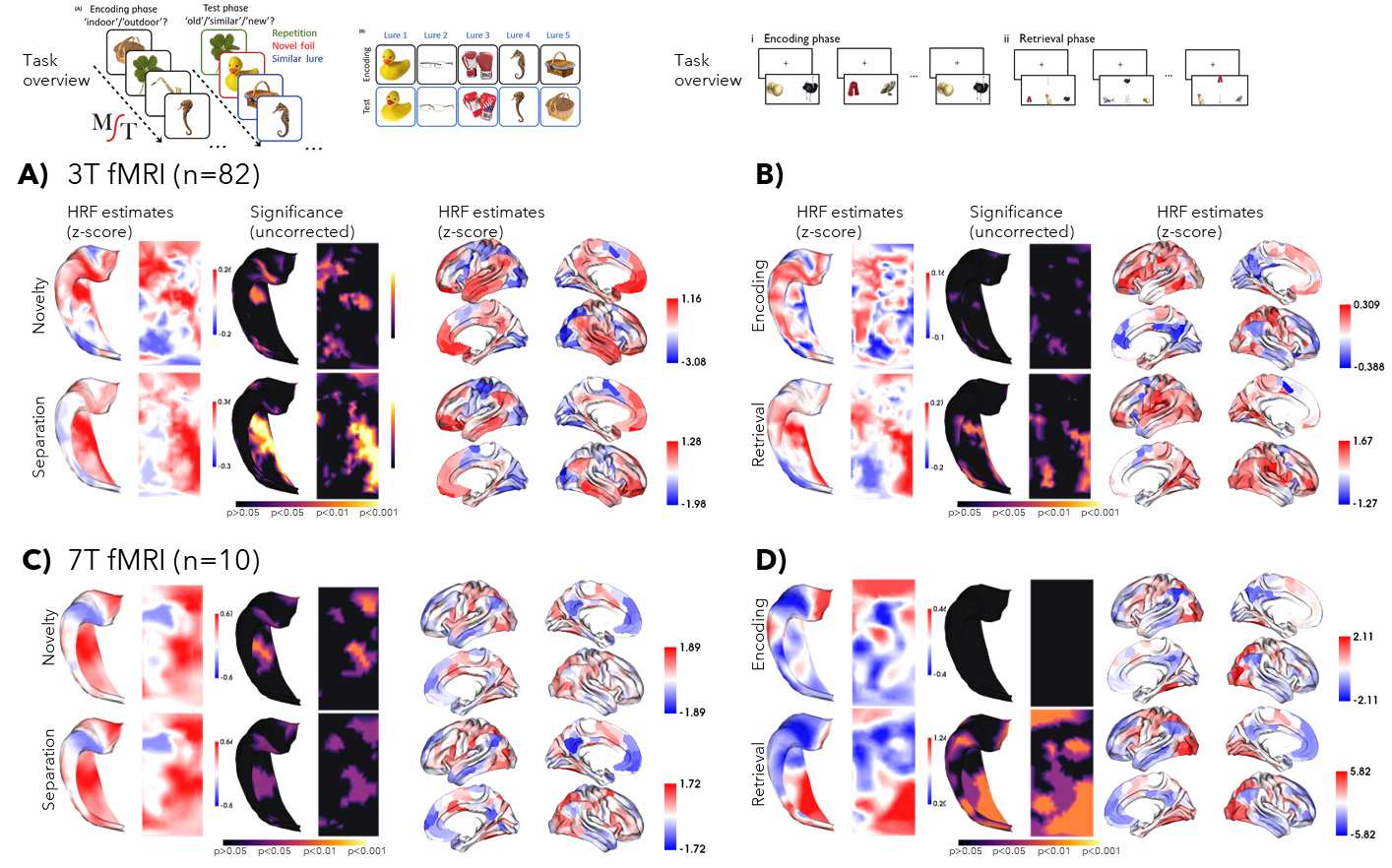

**Figure S7.** Replication of task fMRI analyses from **Figure 7A** at 3T and 7T, and across the MST task and a common object pairing study-test paradigm. **A)** Same data as shown in **Figure 7A**, but with significance testing. **B)** Successfully trials during episodic encoding (subsequent memory paradigm) and retrieval. **C)** and **D)** same as *A)* and *B)*, but performed with 7T fMRI and a smaller cohort.

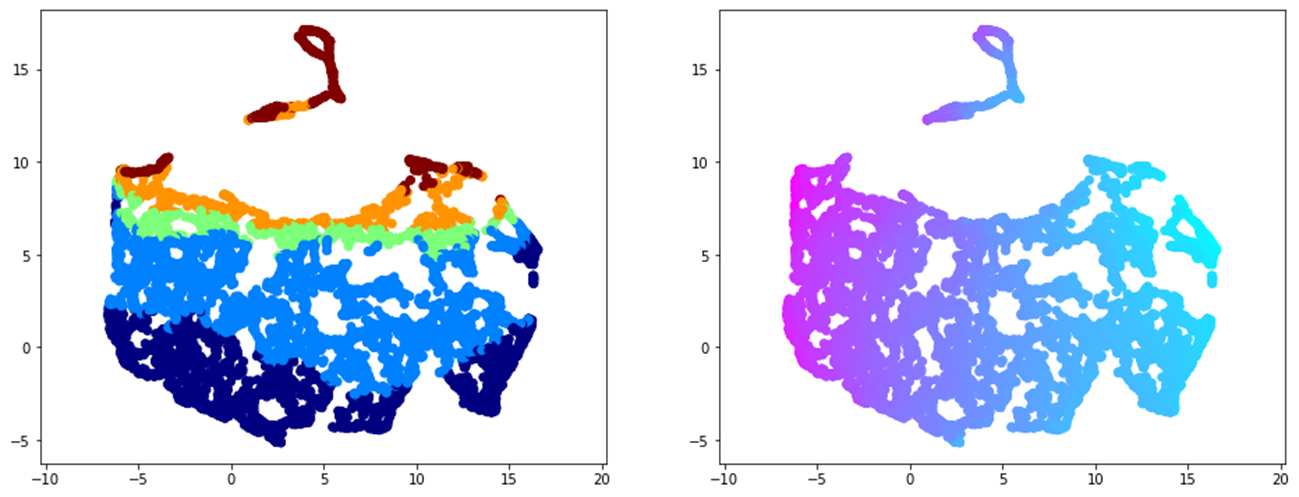

**Figure S8**. Uniform Manifold Approximation and Projection (UMAP) vertices using feature maps from all methods. Coloured according to subfield (left) and anterior-posterior location (right). All default parameters used (McInnes and Healy, 2018).
