## Supplementary Methods for "HippoMaps: multiscale cartography of human hippocampal organization"

**Histology**

BigBrain, both AHEAD brain samples, and one 3D PLI hemisphere were manually segmented and processed with *HippUnfold* in our previous work (DeKraker *et al.*, 2020; DeKraker *et al.*, 2023). Quantitative data were then sampled on hippocampal surfaces using the Connectome Workbench tool “volume-to-surface-mapping” with trilinear interpolation (https://www.humanconnectome.org/software/connectome-workbench). In the current work, we sampled 100μm^3^ images to save computational resources (summarized for all data in Table S1). Masks of local outliers were defined as being more than four standard deviations from the average of a ten vertex radius and were then dilated by two vertices to discard data at an interface with missing areas where excess stain or other artifacts can accumulate. Surface-based interpolation (linear) and extrapolation (nearest-neighbour) were used to fill in missing data. Finally, microstructural profiles over 25 surface depths were aligned using the methods in **Figure 1B**. This was especially important since some of the AHEAD brain data showed misalignment between modalities, leading to some vertices being outside of the hippocampus in some cases, depending on the image. By extrapolating these surfaces beyond the original inner and outer hippocampal bounds, we aimed to capture the actual hippocampal tissue intensities rather than those of the surrounding white matter or background. These were then aligned by translation in the laminar direction that maximized correlation to the global average microstructural profile. Translations were limited to 25% of the surface thickness at each column/vertex.

**Table S1.** Summary of imaging data used in HippoMaps.

| **Dataset** | **Modality** | **Resolution acquired** | **Resolution employed** | **Samples** (x hemispheres) |
| --- | --- | --- | --- | --- |
| BigBrain | Merker | 20μm^3^ | 100μm^3^ | 1(x2) |
| 3D PLI | Transmittance | 48x48x60μm | 100μm^3^ | 1(x1) |
| AHEAD | Bielschowsky  Blockface  Calbindin  Calretinin  Parvalbumin  Thionin  Proton Density  qR1  qR2* | 150x150x200μm | 100μm^3^ | 2(x2) |
| PNI 7T | qT1  T2*  MTR  DWI  rsfMRI  task fMRI | 0.5mm^3^  0.7mm^3^  0.72mm^3^  1.1mm^3^  1.9mm^3^  1.9mm^3^ | 0.25mm^3^  0.7mm^3^  0.72mm^3^  1.1mm^3^  1.9mm^3^  1.9mm^3^ | 10(x2) |
| MICs 3T | qT1  DWI  rsfMRI  task fMRI | 0.8mm^3^  1.6mm^3^  3.0mm^3^  3.0mm^3^ | 0.8mm^3^  1.6mm^3^  3.0mm^3^  3.0mm^3^ | 99(x2)  99(x2)  88(x2)  82(x2) |
| MICs 3T  TLE patients | qT1 | 0.8mm^3^ | 0.8mm^3^ | 33(x1) |

**MRI**

In the PNI (Cabalo *et al.*, 2024) dataset, 10 healthy subjects (5M/5F, age=26.8±4.61 years) underwent three imaging sessions to include additional acquisition modalities and for re-scanning to provide additional image contrast. T1 images were acquired with an MP2RAGE sequence with the following parameters: voxel size 0.5mm^3^, repetition time 5170ms, echo time 2.44ms, flip angle 4^o^, field of view 260x260mm^2^, iPAT acceleration factor 3, partial Fourier 6/8, 320 sagittal slices, with a final matrix size 320x320x320. Superresolution sampling was performed to improve the contrast available in these images. That is, all qT1 images were upsampled from 0.5mm^3^ to 0.25mm^3^, and rigidly registered usings ANTs (Avants *et al.*, 2009) to the first qT1 image for each participant. These high resolution, high contrast images were then run through *HippUnfold* *v1.3.0*. Diffusion weighted images (DWI) were acquired with the following parameters: voxel size 1.1mm^3^, repetition time 7383ms, echo time 70.60ms, flip angle 90^o^, refocusing angle 180^o^, field of view 211x211mm^2^, multiband factor 2, echo spacing 0.79ms, with three shells at b-values 300, 700, and 2000s/mm^2^. b0 images acquired in reverse phase encoding direction are also provided for distortion correction of DWI scans. Derivative apparent diffusivity coefficient (ADC) and fractional anisotropy (FA) were generated within *micapipe*, which employs *MRtrix* (Tournier *et al.*, 2012). Magnetic transfer ratio (MTR) was acquired with the following parameters: voxel size 0.7mm^3^, repetition time 95ms, echo time 3.8ms, flip angle 5^o^, field of view 230x230mm^2^, 240 sagittal slices, pulse width 50ms, and inter-pulse delay 1ms. T2* was acquired with the following parameters: voxel size 0.7mm^3^, repetition time 43ms, multi-echo times 6.46, 11.89, 17.33, 22.76, 28.19, and 33.62ms, flip angle 13^o^, field of view 220x220mm^2^, 160 sagittal slices. Each of these structural images were rigidly registered to this superresolution qT1 image and sampled on hippocampal surfaces provided by *HippUnfold*, as above. Resting state and task fMRI data were acquired using an EPI sequence with the following parameters: voxel size 1.9mm^3^, repetition time 1690ms, multi-echo times 10.80, 27.3, and 43.8ms, flip angle 67^o^, field of view 224x224mm^2^, 775 slices oriented to AC-PC-39^o^, multiband factor 3, and echo spacing 0.53ms.

The MICs dataset release initially included 50 subjects (Royer *et al.*, 2022), but here we include proceeding scans with the same protocol from 99 healthy subjects (54M/45F, age=29.6±9.7 years) and 33 temporal lobe epilepsy (TLE) patients (12M/11F, age=42.4±11.0 years). Patient scans were compared to only a subset of the MICs data to maximize demographic similarity (21M/21F, age=40.4±10.4 years). qT1 scans were acquired with an MP2RAGE sequence with the following parameters: voxel size 0.8mm^3^, repetition time 5000ms, echo time 2.9ms, flip angle 5°, iPAT acceleration factor 3, bandwidth = 270 Hz/px, partial Fourier  6/8, 240 sagittal slices, with a final matrix size of 320x320x240. Diffusion weighted images (DWI) were acquired with the following parameters: voxel size 1.6mm^3^, repetition time 3500ms, echo time 64.40ms, flip angle 90^o^, refocusing angle 180^o^, field of view 240x240mm^2^, multiband factor 3, echo spacing 0.76ms, b-value 0, with three shells at b-values 300, 700, and 2000s/mm^2^. b0 images acquired in reverse phase encoding direction are also provided for distortion correction of DWI scans. Resting state fMRI data were acquired using an EPI sequence with following parameters: voxel size 3.0mm^3^, repetition time 600ms, echo time 30ms, flip angle 52^o^, field of view 240x240mm^2^, multiband factor 6, and echo spacing 0.54ms. All scans were preprocessed using *micapipe v0.2.0* (Cruces *et al.*, 2022) followed by *HippUnfold* *v1.3.0*. Each of these structural images was registered within *micapipe* and sampled on hippocampal surfaces provided by *HippUnfold*, as above.

Functional MRI data were sampled onto hippocampal surfaces with an average vertex spacing of 2mm, in correspondence with the lower resolution of this data, making 419 vertices x N timepoints. Custom code was used to compute intrinsic timescale, regional homogeneity, and functional connectivity analyses of resting state data. For task data, nilearn (Huntenburg *et al.*, 2017) was used to construct general linear models (GLMs). All trial types were modeled separately after convolution with a haemodynamic response function and its derivative and dispersion, alongside regressors of no interest for motion (6 degrees of freedom) and drift (8 frequencies).

For comparison between TLE patients and healthy controls, a subset of 81 controls was selected to match age and sex of the TLE group. All TLE patients had a clear lateralization of primary epileptogenic zones (15L/20R). Only ipsilateral hemispheres were examined here.

**iEEG**

Locally collected data came from 20 epilepsy patients (7M/13F; age=33.90±9.02 years) implanted with deep intracranial electrodes as part of their standard of care. Only channels within the brain, free of artifacts, no ictal or interictal events, and during resting wakefulness with eyes closed were examined, making up 2507 channels across the whole brain. When no continuous and artifact-free 60-second segments were found for a given channel, multiple discontinuous segments from the channel were concatenated with a 2-second zero-padded buffer between segments. Each patient also had an intraoperative CT or MRI scan, from which all channels were located using the Intraoperative Brain Imaging System (Ibis) (Drouin *et al.*, 2017). These scans were then rigidly registered to preoperative T1w images that were run through *HippUnfold*. From all channels, 44 (7 left; 37 right hemisphere) were within 5mm of any hippocampal vertex. These were concatenated with similarly preprocessed data from (Frauscher *et al.*, 2018) consisting of 1772 channels in standardized MNI152 (2009c, non-linear, symmetric) volumetric space. *HippUnfold* was run on the MNI152 T1w template (https://github.com/khanlab/hippunfold-templateflow), allowing us to localize hippocampal electrodes (albeit, with less subject-specific precision). From all channels 37 (19 left; 18 right hemisphere) were within 5mm of any hippocampal vertex. Power spectrum densities were then calculated for the combined datasets using Welch’s method with a Hann window. Power spectrum densities were then linearly resampled to a range of 0.5 to 80Hz. Band powers for delta, theta, alpha, beta, and gamma bands were calculated in the range of 0.5-4Hz, 4-8Hz, 8-13Hx, 13-30Hz, and 30-80Hz, respectively. Extrapolation of these sparsely sampled hippocampal vertex data was then performed using the method shown in **Figure 1C**. That is, typical nearest neighbour or linear interpolations were inappropriate since they would only consider one or two closest channels. Thus, we used a method that averages channels proportionally to their geodesic distance to a given vertex to maintain robustness while still providing some level of spatial differentiation based on where a given channel originated from.

Full details and code for each method described above are available within their respective online tutorials, while have been made simple and well commented to serve as examples for future hippocampal mapping projects at https://github.com/MICA-MNI/hippomaps.

**Microstructural profile alignment**

Microstructural profiles are highly sensitive to errors in tissue segmentation which can cause a profile to shift in the horizontal direction (that is, towards the superficial or deep parts of the cortex). We aimed to account for this sensitivity by including an additional post-processing step to vertically translate profiles until they are maximally aligned. This was accomplished by first sampling laminar depths both within and outside (above and below) the hippocampal cortical mantle to collect data from any tissue that may have been under segmented. We then translate these profiles vertically by shifting them up or down by a given number of depth-wise surfaces. The amount of translation is that which maximizes the alignment of a given profile to the global average of all profiles from a given map (that is, across all vertices). This is shown in an example in **Supplementary Methods Figure S1**.

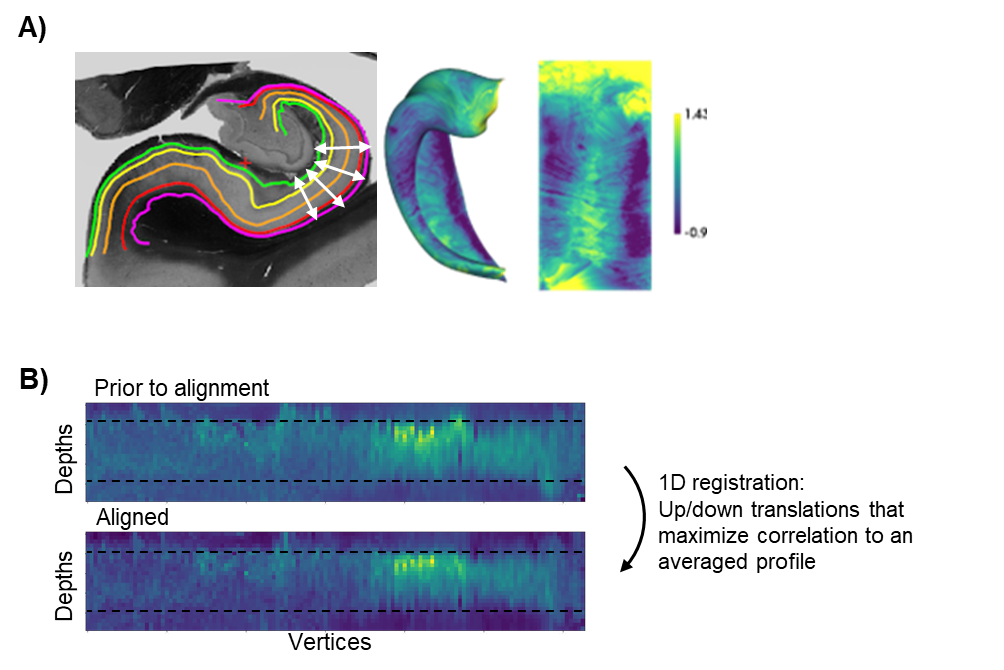

**Figure S1.** Alignment of microstructural profiles by vertical translations. **A)** One slice from 3D PLI data illustrating surfaces at various depths through the cortical mantle (yellow, orange, and red), as well as surfaces extrapolated above and below the cortical mantle (green and violet). **B)** Profiles across equivalent vertices from all surfaces, shown for a set of 500 vertices. The segmented cortical mantle boundaries are indicated by the dotted lines. Image intensities more closely follow these boundaries after alignment.

**Permutation testing**

The HippoMaps tools include several methods for permutation testing of spatial correlations, most notable Spin test correction (Karat *et al.*, 2023), Moran spectral randomization (Wagner & Dray, 2015), and Eigenstrapping (Koussis *et al.*, 2024). These methods are overviewed in **Figure S2**, including an example of what a random permutation can look like. The Spin test was observed to be most conservative, while Moran permutation was least conservative (that is, most likely to show a false positive as in the example shown).

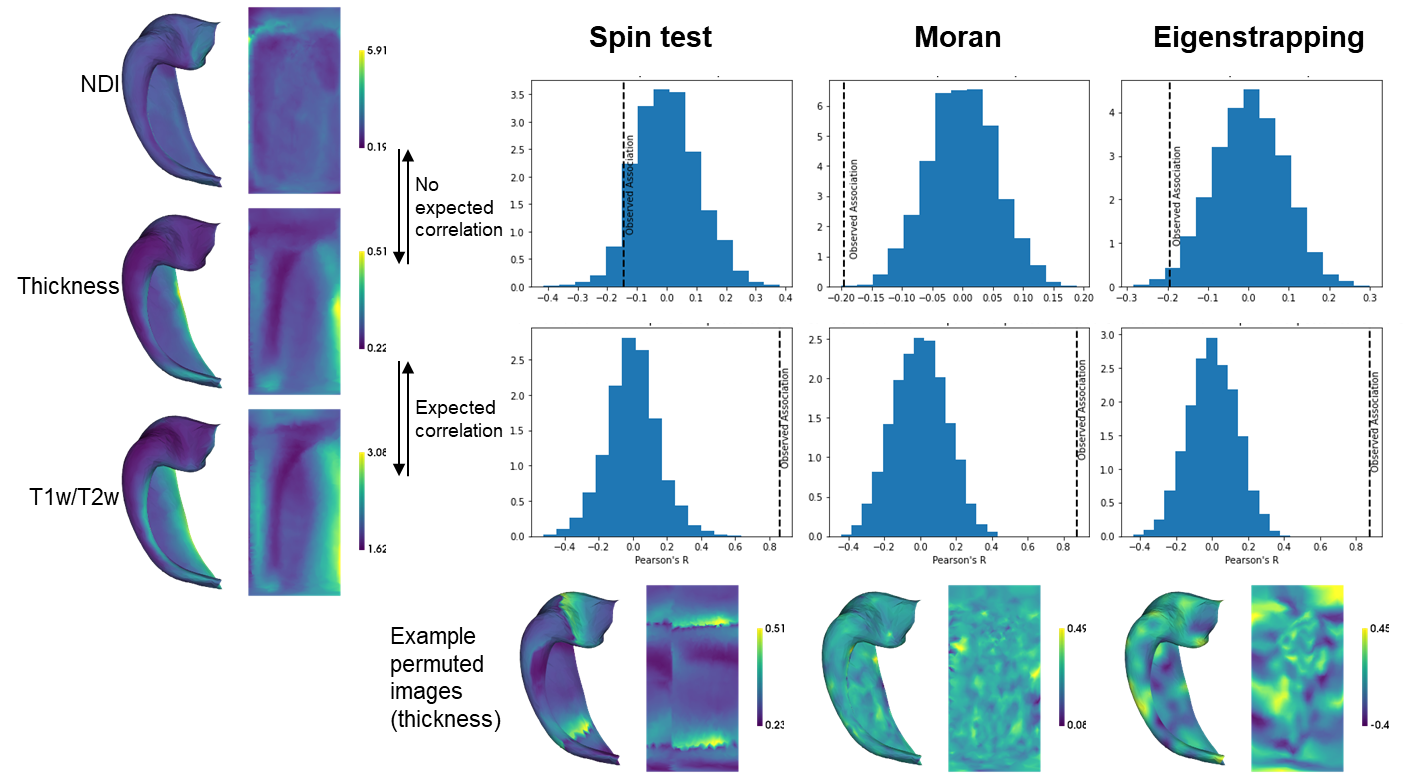

**Figure S2.** Permutation tests for spatial correlation of hippocampal maps. Example Neurite Density Index (NDI), thickness, and T1w over T2w ratios are shown on the left, with data from (Karat *et al.*, 2023). To the right, a null distribution of correlation with permuted data is shown for each method. The observed R value is indicated by the dotted line. If the observed R value is within the null distribution, then there is no significant correlation. The p-value can be calculated as the proportion of the null distribution greater than the observed R value. One example of a permuted thickness map is shown below for each method.
